# *Pseudomonas aeruginosa* TfpW is a multifunctional D-Ara*f* glycosyltransferase and oligosaccharyltransferase

**DOI:** 10.1101/2020.04.19.049825

**Authors:** Anne D. Villela, Hanjeong Harvey, Katherine Graham, Lori L. Burrows

## Abstract

TfpW is an oligosaccharyltransferase that modifies the subunits of type IV pili from group IV strains of *Pseudomonas aeruginosa* with oligomers of α-1,5-linked-D-arabinofuranose (D-Ara*f*). Besides its oligosaccharyltransferase activity, TfpW may be responsible for periplasmic translocation and polymerization of D-Ara*f*. Here we investigated these potential roles of TfpW in Pa5196 pilin glycosylation. Topology studies confirmed the periplasmic location of loop 1 and the large C-terminus domain, however the central portion of TfpW had an indeterminate configuration. Reconstitution of the Pa5196 pilin glycosylation system by providing *pilA*, *tfpW* +/- *tfpX* and the D-Ara*f* biosynthesis genes *PsPA7_6246-6249* showed that TfpW is sufficient for glycan polymerization and transfer to pilins in *P. aeruginosa* PAO1, while TfpX is also necessary in *Escherichia coli*. In addition to PsPA7*_*6246, DprE1 (PsPA7*_*6248) and DprE2 (PsPA7*_*6249), the GtrA-like component PsPA7*_*6247 was required for pilin glycosylation in *E. coli* versus PAO1. In a PAO1 Δ*arnE/F* mutant, loss of PsPA7_6247 expression decreased the level of pilin glycosylation, suggesting that *arnE/F* may play a role in pilin glycosylation when PsPA7_6247 is absent. Bacterial two-hybrid studies showed interactions of TfpW with itself, TfpX, PsPA7*_*6247 and DprE2, suggesting the formation of a complex that enables efficient pilin glycosylation. Fluorescence microscopy of *E. coli* and Pa5196*ΔdprE1* expressing a DprE1-sGFP fusion showed that the protein is expressed in the cytoplasm, supporting our model that includes cytoplasmic biosynthesis of the lipid carrier-linked D-Ara*f* precursor prior to its periplasmic translocation. Together these data suggest that TfpW may be the first example of a trifunctional flippase, glycosyltransferase, and oligosaccharyltransferase.

## INTRODUCTION

*Pseudomonas aeruginosa* is an opportunistic pathogen and the major cause of morbidity and mortality among cystic fibrosis patients (1). It is among the pathogens listed as ‘critical’ by the World Health Organization for new antibiotic development. Among the important virulence factors of *P. aeruginosa* are its type IV pili (T4P) (2,3). T4P are expressed on the cell surface and are involved in adherence, motility, and pathogenesis (2,3). They are helical protein polymers containing thousands of copies of the major pilin, PilA. Five different PilA variants (groups I through V) were identified in *P. aeruginosa* strains, differing in amino acid sequence, association with unique accessory genes, and post-translational modifications (4). Pilins from group IV (e.g. strains Pa5196 and PA7), are modified with homo-oligomers of α-1,5-linked-D-arabinofuranose (D-Ara*f*), O-linked to multiple Ser and Thr residues, predominately at Thr64 and Thr66 (5,6). The same D-Ara*f* configuration is found in arabinogalactan (7) and lipoarabinomannan (8), major components of the *Mycobacterium tuberculosis* cell wall. Antisera against LAM from *M. tuberculosis* cross-reacts with D-Ara*f* modified pilins, while Pa5196 pilin antisera recognizes cell wall material from *Mycobacterium smegmatis* (5).

Pilin glycosylation requires synthesis of the glycan, its translocation to the periplasm, and transfer to the protein acceptor. The *P. aeruginosa* biosynthetic pathway for D-Ara*f* is encoded by a cluster of seven genes (*PsPA7*_*6245-6251;* the *PsPA7* prefix will be omitted below for brevity) (9). 6245, 6246, 6248 and 6249 are homologues of the *M. tuberculosis* H37Rv genes Rv3807c (encoding decaprenyl-phosphoryl-5-phosphoribose phosphatase), Rv3806c (encoding decaprenyl-P-ribose-5-P, DPPR, synthetase), Rv3790 (encoding decaprenylphosphoryl-β-D-ribose oxidase, DprE1) and Rv3791 (encoding decaprenylphosphoryl-2-keto-β-D-erythro-pentose reductase, DprE2) (9). The remaining genes 6247, 6250, and 6251 encode hypothetical proteins. Expression of 6246, DprE1, and DprE2 is essential for Pa5196 twitching motility and pilin glycosylation (9). A Pa5196 6245 mutant produced less heavily glycosylated pilins, while loss of 6250 and 6251 expression did not alter glycosylation (9). No homologues of *emb*, encoding the *Corynebacterium* enzyme that generates α-1,5-linked D-Ara*f* polymers, were identified in the *P. aeruginosa* D-Ara*f* biosynthetic cluster (9). The mechanism by which D-Ara*f* is translocated to the periplasm, either as single sugars or as α-1,5-linked oligosaccharides, is also unknown.

TfpW is a putative glycosyltransferase (GT) belonging to a family of membrane-bound GT-C enzymes and has very limited amino acid sequence similarity to *Corynebacterium* Emb and related arabinosyltransferases such as *M. tuberculosis* EmbA/B/C (5). *tfpW* is immediately downstream of *pilA* in strains Pa5196 and PA7, and encodes a 646 amino acid product with a cytoplasmic N-terminus, 11 predicted transmembrane α-helices and a large periplasmic C-terminal region containing a tetratricopeptide repeat (TPR) structural domain (5). TPR motifs consist of ∼34 residue helical hairpins arranged as tandem repeats. They mediate protein-protein or protein-sugar interactions, and assembly of multiprotein complexes (10). Loss of TfpW abrogates pilin glycosylation, including the addition of single sugars (5). Complementation of a *tfpW* mutant with TfpW containing mutations in the conserved glycosyltransferase motif (DDX) found in GT-C superfamily members failed to restore pilin glycosylation, but it was not clear whether that was due to loss of oligosaccharide biosynthesis or of glycan transfer to the protein acceptor (5). The *tfpX* gene, located downstream of *tfpW*, cannot mediate glycosylation, since complementation of a Pa5196 *tfpW* mutant with *pilA*+*tfpX* failed to restore modification (5).

Phylogenetically, the *P. aeruginosa* group IV pilin is more similar to PilE from *Neisseria gonorrhoeae* and *N. meningitidis* than to other *P. aeruginosa* pilins (4). In *Neisseria*, cytoplasmic enzymes generate undecaprenol-linked oligosaccharides and the PglF flippase translocates them to the periplasm where they are attached to pilins by the PglL O-oligosaccharyltransferase (11). PglL has extreme substrate promiscuity, transferring any glycan from the lipid carrier to pilins (12). In group I strains of *P. aeruginosa*, the oligosaccharyltransferase TfpO (PilO) has a relaxed glycan specificity (13), transferring a lipopolysaccharide (LPS) O-antigen unit (3-5 sugars, depending on the serotype) to the C-terminal Ser of cognate pilins. In that case, undecaprenol-linked O-antigen units are first synthesized by the cytoplasmic LPS biosynthetic machinery, then translocated to the periplasm by the O-unit flippase Wzx (3), where they are accessed by TfpO.

No flippase for the D-Ara*f* glycans that decorate group IV pilins has been identified. Although it lacks sequence similarity with PglF or Wzx, TfpW’s predicted topology is compatible with that of flippases. It could be responsible for the translocation of lipid-linked D-Ara*f* precursors, either single sugars or α-1,5 linked oligosaccharides. Further, TfpW’s weak similarity to Emb enzymes and essential DDX motif suggest it has GT activity, consistent with forming α-1,5 linked D-Ara*f* oligosaccharides. Therefore, TfpW may be a flippase, glycosyltransferase, and oligosaccharyltranferase.

Here we further investigated the potential roles of TfpW in Pa5196 pilin glycosylation, important for pilin folding, stability, assembly, and function (5,9). Topology and structure prediction studies were used to evaluate conserved regions of TfpW that may be responsible for its potential activities. The Pa5196 pilin glycosylation system was reconstituted in PAO1, a laboratory strain that expresses non-glycosylated pili, showing that TfpW was both necessary and sufficient for polymerization of the glycan and its transfer to pilins. Parallel experiments in *Escherichia coli* showed that additional components – including TfpX – were needed for pilin glycosylation in that background. Interactions of TfpW with itself, TfpX, and proteins involved in D-Ara*f* biosynthesis suggest they form a glycosylation complex for efficient pilin modification.

## RESULTS

### tfpW, but not tfpX, is conserved in T4P-encoding species

We observed previously that deletion of *tfpX* caused pilin glycosylation defects. However, bioinformatic analyses showed that while *tfpW* is encoded with *pilA* in multiple species, *tfpX* is rare (Fig. 1A). These data support the hypothesis that TfpW is sufficient for pilin glycosylation. In species such as *Pseudomonas indica* and *Kingella denitrificans*, the *pilA* and *tfpW* genes are clustered with the D-Ara*f* biosynthetic genes. Putative TfpW homologues from *Azotobacter beijerinckii*, *Acinetobacter venetianus*, *Teredinibacter turnerae*, *P. indica* and *K. denitrificans* have 29.9 to 47.1% identity to TfpW from *P. aeruginosa* PA7 (Fig. 1B). The regions conserved among putative TfpW homologs are located next to or in periplasmic loops (14) 1, 2, 3, 4 and 5 as predicted by CCTOP (Constrained Consensus TOPology prediction server) (Fig. 1B). Although the sequence similarity of the C-terminal regions is low, Phyre2 analyses predict that all contain TPR domains (15) (data not shown).

**Figure 1.**
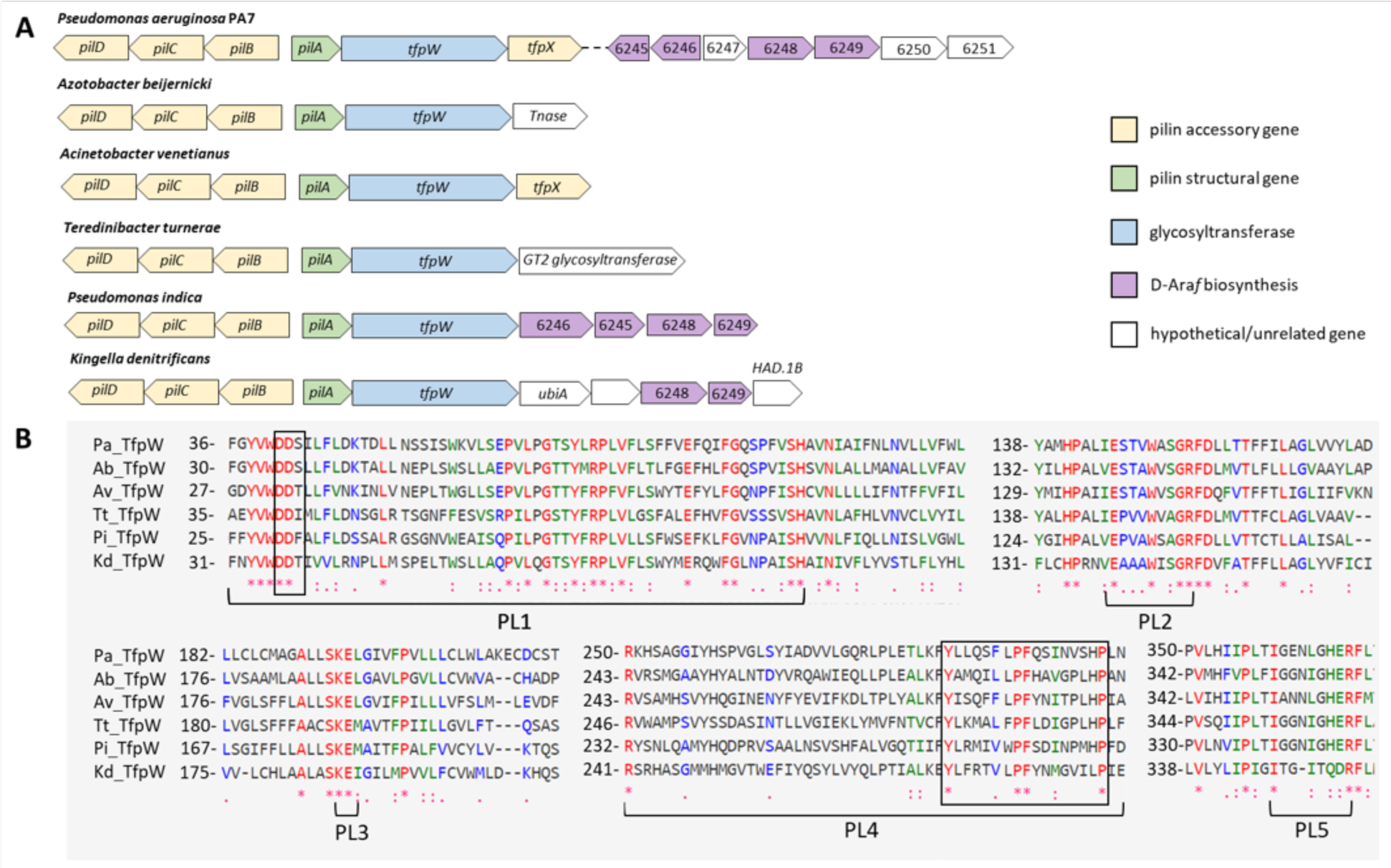
Genomic organization of *tfpW* from *P. aeruginosa* PA7 and putative homologs from different bacterial species (A), and selected regions of amino acid sequence alignment (B). **A.** The *tfpW* is located downstream to *pilA* in all bacterial species, but *tfpX* is not. In *P. indica* and *K. denitrificans* the genes encoding the biosynthesis of D-Ara*f* are clustered with the *pilA* and *tfpW* genes. **B.** The conserved regions among the putative TfpW homologs are located next to or on the predicted periplasmic loops (14) 1, 2, 3, 4 and 5. The conserved DDX domain is located on PL1, while the proline-rich domain (YX_6_PFX_7_P) is located on PL4 (boxed). Pa, *Pseudomonas aeruginosa* PA7 (GenBank accession number AAM52060.1); Ab, *Azotobacter beijerinckii* (WP_090729738.1); Av, *Acinetobacter venetianus* (WP_130134296.1); Tt, *Teredinibacter turnerae* (WP_028883051.1); Pi, *Pseudomonas indica* (WP_139198627.1); Kd, *Kingella denitrificans* (STR11763.1). Position of the first amino acid in the alignment is indicated before each selected sequence.

### TfpW contains a proline-rich motif conserved in Emb proteins

*P. aeruginosa* TfpW has limited amino-acid sequence similarity with other oligosaccharyltransferases such as TfpO or PglL, all of which have multiple transmembrane segments followed by a large C-terminal periplasmic domain. In *P. aeruginosa* group I and group IV strains, the genes encoding TfpO and TfpW are located immediately downstream of *pilA* encoding the major pilin, while in *N. meningitidis, pglL* and *pilE* (the equivalent of *pilA*) are unlinked (Fig. 2A). In *M. tuberculosis* genes the encoding glycosyltransferases AftA and EmbA/B/C that synthesize D-Ara*f-*containing glycans are clustered with genes involved in D-Ara*f* precursor biosynthesis (Fig. 2A). No *emb*-like genes were identified in *P. aeruginosa*, and the presence of a DDX motif characteristic of GT-C family enzymes in TfpW suggested that it could have glycosyltransferase (forming the α-1,5 linkages between D-Ara*f* monomers) and oligosaccharyltransferase (transferring D-Ara*f* oligosaccharides to the pilin acceptor) activities. Despite TfpW’s limited sequence identity with Emb proteins, a second motif similar to the proline-rich domain in *M. tuberculosis* EmbC (YX_6_PX_5_P) (16) was identified in putative PL4, near the C-terminal TPR domain (Fig. 2B). This motif is found in membrane-bound polysaccharide polymerases and regulates polysaccharide chain length (16,17). The YX_6_PFX_7_P sequence was conserved among putative TfpW homologs (Fig. 1B).

**Figure 2.**
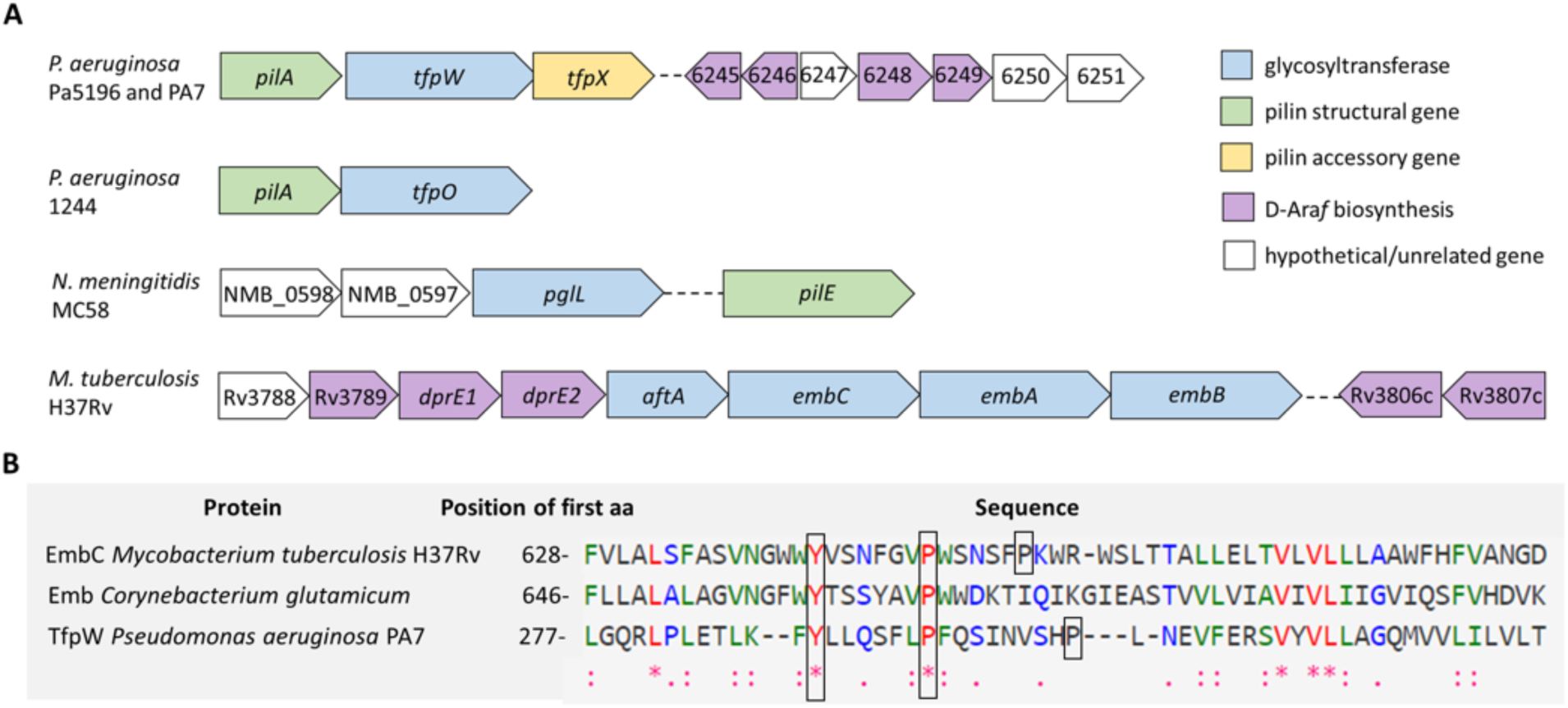
Genomic organization of pilin glycosyltransferases-encoded genes in *P. aeruginosa* PA7, 1244, *N. meningitidis* and arabinosyltransferases-encoded genes in *M. tuberculosis* H37Rv (A), and selected region of amino acid sequence alignment of EmbC from *M. tuberculosis* H37Rv, Emb from *Corynebacterium glutamicum* and TfpW (B). **A.** In *P. aeruginosa* PA7 and 1244, the genes that encode glycosyltransferases *tfpW* and *tfpO*, respectively, are located downstream of *pilA*, while in *N. meningitidis pglL* and *pilE* are unlinked. In *M. tuberculosis* the genes that encode for glycosyltransferases *aftA* and *embCAB* are clustered with a subset of the genes involved in D-Ara*f* biosynthesis. **B.** A motif similar to the proline-rich domain found in EmbC from *M. tuberculosis* (YX_6_PX_5_P) is present in TfpW (YX_6_PFX_7_P).

### Topology of TfpW

TfpW has 11 predicted transmembrane segments, with a cytoplasmic N-terminus and periplasmic C-terminus. In the absence of a structure, we probed its topology experimentally using dual reporter (PhoA/LacZα) fusions at select positions (8). Periplasmic fusions result in blue colonies due to alkaline phosphatase activity, while cytoplasmic fusions yield red colonies due to β-galactosidase activity. Transmembrane or indeterminate fusions result in purple colonies. Only fusions in the first predicted periplasmic loop and in the large C-terminal domain gave a strong alkaline phosphatase signal (Fig. 3A), while fusions to other putative cytoplasmic and periplasmic loops resulted in purple colonies (Fig. 3A). One fusion predicted to be located in the membrane (G245) instead resulted in blue colonies, suggesting a periplasmic localization. These data suggest that the configuration of the central portion of the protein does not match *in silico* predictions. We generated a structural model of TfpW using the Phyre2 server (Fig. 3B **and** C). The closest structural template was ArnT, a 4-deoxy-4-aminoarabinose transferase from *Cupriavidus metallidurans*. Most of TfpW was modelled with high confidence up to the C-terminal TPR domain, while the best independent match to TfpW’s TPR domain was the C-terminal domain of human O-GlcNAc transferase. Putative TM segments 5 through 8 in the ArnT-based model were distorted, especially in the 180° view (Fig. 3C). This result, coupled with our topology data, suggests that the central portion of TfpW has an unusual fold.

**Figure 3.**
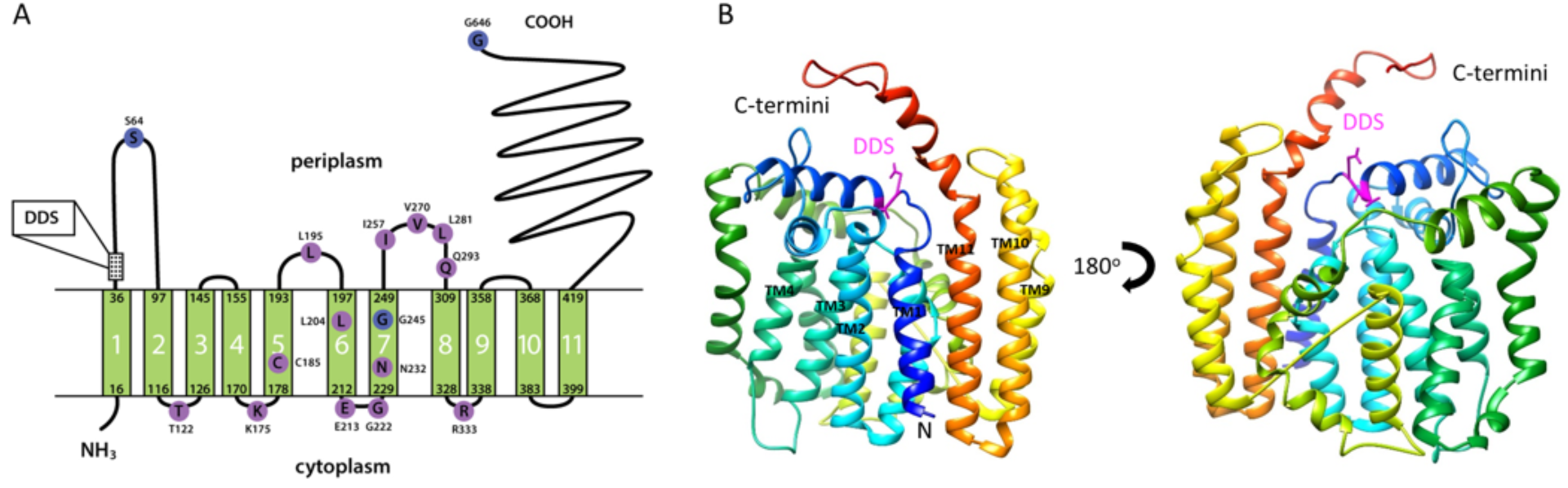
TfpW topology and structural prediction. **A.** TfpW is predicted to have a cytoplasmic N-terminus, 11 transmembrane (TM) segments, 5 periplasmic loops and a large periplasmic C-terminal domain (CCTOP). Dual reporter assay to evaluate topology of TfpW in *E. coli* DH5α strain. Fusion of PhoA/LacZα with amino acids localized in periplasmic region result in blue colonies due to alkaline phosphatase activity (blue cycles), while fusion with cytoplasmic amino acids lead to obtainment of red colonies due to β-galactosidase activity. Purple colonies are observed when fusion is in transmembrane region (purple cycles). Loops in the central region of TfpW have ambiguous topology. Dual indicator plates contained ampicillin (100 µg/mL), IPTG (1 mM), BCIP (80 µg/mL) and Red-Gal (100 µg/mL). **B.** Structure prediction of TfpW, based on the ArnT template, a 4-deoxy-4-aminoarabinose transferase from *Cupravidas metallireducans* (Phyre 2,(15)). TM domains 5, 6, 7 and 8 in this model seem unreliable, especially in the 180° view.

### PilA_5196_ glycosylation can be reconstituted in PAO1 and E. coli

To establish whether *pilA, tfpW*, and the D-Ara*f* biosynthetic genes are sufficient for PilA_5196_ glycosylation in a heterologous system, we introduced constructs encoding various subsets of these genes into *P. aeruginosa* PAO1 (which lacks the genes for D-Ara*f* biosynthesis) as well as into *Escherichia coli*. We previously found that loss of *tfpX* prevented pilin glycosylation (5), but its lack of conservation in other species with *tfpW* homologues suggested that it may be dispensible. As *pilA*, *tfpW* and *tfpX* are co-transcribed, we wondered if deletion of *tfpX* was detrimental to transcript stability. We generated *pilA-tfpWX* constructs with a frameshift mutation at Glu16 of TfpX (expressing only first 15 amino acids) or a stop codon at Leu56. In both cases, pilins were still modified in the PAO1 background (Fig. 4A), suggesting that TfpX is not required. Instead, deletion of *tfpX* (AW, Fig. 4A) likely disrupts TfpW expression. Interestingly, TfpX expression was necessary for pilin glycosylation in *E. coli*, since no modification was observed using the construct with a frameshift mutation at Glu16 (Fig. 4B). *E. coli* lacks the machinery for surface assembly of *P. aeruginosa* pilins, but we could still monitor pilin expression and post-translational modification by Western blot of cell lysates using specific antibodies. In addition to TfpX, we found that 6247 – a hypothetical protein that shares ∼59-63% sequence identity with GtrA proteins from other bacteria – plus 6246, DprE1 and DprE2 were required for PilA_5196_ glycosylation in *E. coli* (Fig. 4C). Members of the GtrA family are involved in the synthesis of cell surface polysaccharides (18).

**Figure 4.**
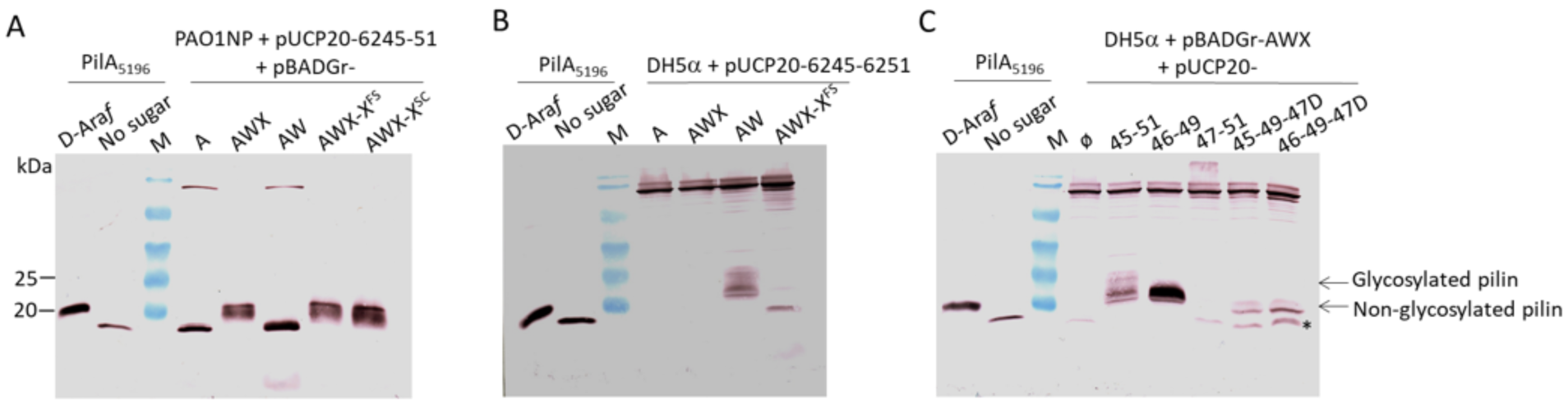
Reconstitution of Pa5196 pilin arabinosylation in PAO1 (A) and *E. coli* (B and C). Cell lysates were prepared from cultures grown in LB agar containing 0.05% L-arabinose, 200 µg ml^−1^ carbenicillin, 30 µg ml^−1^ gentamicin for PAO1 or 0.2 % L-arabinose, 100 µg ml^−1^ ampicillin, 15 µg ml^−1^ gentamicin for *E. coli*. Western blot was performed with anti-PilA_IV_ primary antibody. **A.** In the PAO1-NP pUCP20-6245-6251 background, PilA_5196_ is glycosylated when *tfpW* is expressed even when *tfpX* has a frameshift mutation at Glu16 (expressing only first 15 amino acids, AWX-X^FS^) or a stop codon at the position of L56 (AWX-X^SC^). AWX, AWX-X^FS^ and AWX-X^SC^ were diluted 10X. **B.** In the *E. coli* DH5α pUCP20-6245-6251 background, PilA_5196_ is glycosylated only when both *tfpW* and *tfpX* are expressed, as a frame shift mutation in *tfpX* at Glu16 (AWX-X^FS^) results in unmodified pilins. **C.** PilA_5196_ glycosylation in *E. coli* requires expression of 6246, 6247, DprE1 (6248) and DprE2 (6249), and the AWX cassette. *Lowest molecular weight band is not unmodified pilins.

GtrA-like component 6247 has three predicted transmembrane domains, a periplasmic N-terminus, cytoplasmic C-terminus, and pI of 9.24 (ExPASy Bioinformatic Resource Portal). A similar configuration and isoelectric point are predicted for GtrA family member Rv3789 from *M. tuberculosis* (18) despite their lack of sequence similarity.

### ArnE/F, an undecaprenyl phosphate aminoarabinose flippase may participate in pilin glycosylation in P. aeruginosa when 6247 is absent

A study in *M. smegmatis* ΔRv3789 strains showed that the reduced arabinose content of AG and LAM and accumulation of decaprenyl-monophosphoryl-β-D-arabinose (DPA) were restored upon complementation with the putative undecaprenyl phosphate aminoarabinose flippase *arnE/F* genes from *E. coli*, suggesting that both Rv3789 and ArnE (PmrM)/ArnF (PmrL) promote D-Ara*f* translocation (19). To investigate whether ArnE/F can function in *P. aeruginosa* pilin glycosylation, a PAO1 Δ*arnE/F* mutant was constructed and complemented with pBADGr-AWX plus pUCP20-Ø (empty vector), or pUCP20 with 6245-6251, 6246-6249, or 6246-6249 containing a 6247 deletion. Western blot of whole cell lysates was performed using a PilA_5196_ antibody. Deletion of *arnE/F* had no effect on pilin glycosylation when 6245-6251 were expressed (Fig. 5A **and** B). AWX plus 6246, 6248 and 6249 are the minimum requirements for pilin glycosylation in PAO1 (Fig. 5A), as observed previously (9). However, loss of 6247 expression in PAO1 Δ*arnE/F* decreased pilin glycosylation (Fig. 5A **and** B), suggesting that ArnE/F may participate in pilin glycosylation when 6247 is absent and confirming the results seen in *M. tuberculosis*.

**Figure 5.**
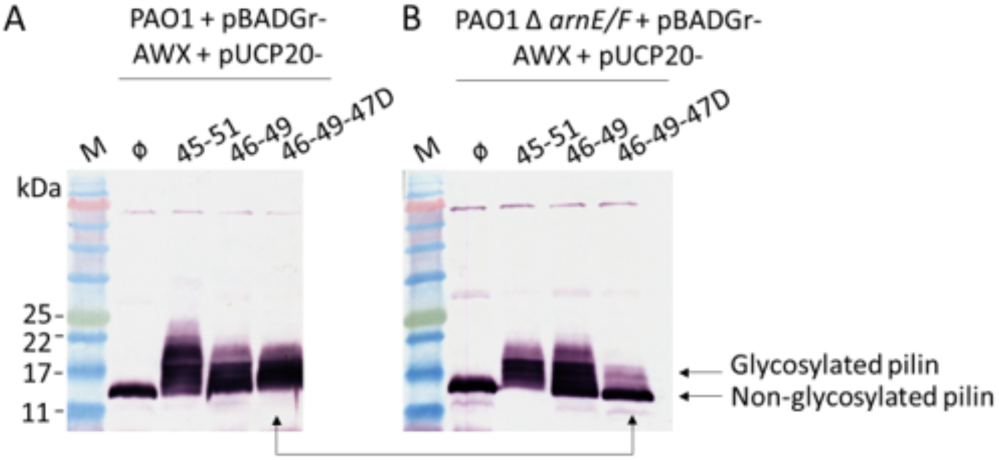
ArnE/F, an undecaprenyl phosphate aminoarabinose flippase, is necessary for pilin glycosylation in *P. aeruginosa* when 6247 is absent. PAO1 (parental) (A) and PAO1 Δ*arnE/F* (B) strains were transformed with pBADGr-AWX and pUCP20-Ø (empty plasmid) / 6245-6251 / 6246-6249/ 6246-6249-Δ6247. Cell lysates were prepared from cultures grown in LB agar containing 0.05% L-arabinose, 200 µg ml^−1^ carbenicillin and 30 µg ml^−1^ gentamicin. Western blot was performed with anti-PilA_IV_ primary antibody.

### TfpW interacts with itself, TfpX and proteins involved in D-Araf biosynthesis

In mycobacteria genes encoding DPA synthesis are clustered with those encoding the Emb glycosyltransferases, while in *P. aeruginosa* the pilin and sugar biosynthetic gene clusters are unlinked. We asked whether the products of the two *P. aeruginosa* gene clusters could form a protein complex to promote efficient pilin glycosylation. Potential interactions were tested using a bacterial adenylate cyclase two-hybrid (BACTH) assay (20). T18-TfpW, -TfpX, -6247, -DprE1 and -DprE2 C-terminal fusions were co-expressed in *E. coli* BTH101 with T25-TfpW, -TfpX, -6247, -DprE1 and -DprE2 C-terminal fusions, and potential interactions were monitored on LB agar containing X-Gal or MacConkey agar containing maltose. TfpW interacted with itself, TfpX, 6247, and DprE2, while 6247 interacted with itself, TfpX, DprE1, and DprE2 (Fig. 6). Interaction of DprE1 and DprE2, previously demonstrated in mycobacteria (21), was also confirmed (Fig. 6). These data suggest that despite being encoded in separate operons, the pilin glycosylation proteins may form a complex.

**Figure 6.**
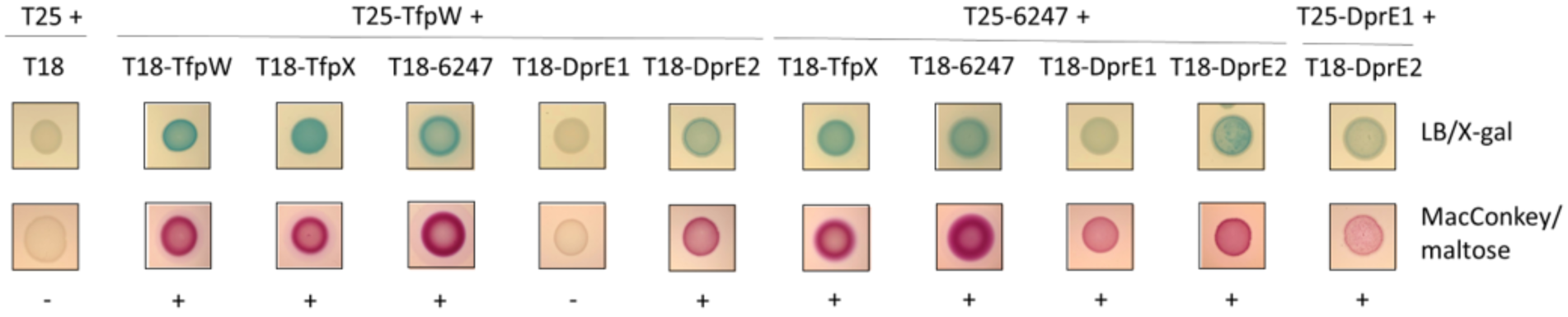
TfpW interacts with itself, TfpX, and with proteins involved in D-Ara*f* biosynthesis. TfpW, TfpX, 6247, DprE1 (6248) and DprE2 (6249) fusions with T18 and T25 fragments of adenylate cyclase were transformed into *E. coli* BTH101 and screened on LB agar/X-gal and MacConkey agar/maltose. TfpW interacts with itself, TfpX, 6247 and DprE2, while 6247 interacts with itself, TfpX, DprE1 and DprE2. Interaction of DprE1 and 2 is also observed. Representative images are shown with positive (+) and negative (-) interactions indicated. At least 3 replicates were performed.

### P. aeruginosa Pa5196 DprE1 is cytoplasmic

A recent study suggested that DprE1 is a highly-accessible drug target due its localization to the periplasmic face of the cytoplasmic membrane of *M. tuberculosis* (22). Periplasmic DPA biosynthesis would preclude the need for a translocase (flippase). The DprE1 proteins of *M. tuberculosis* and *P. aeruginosa* share 33% identity, but neither has a secretion signal, making us skeptical of a non-cytoplasmic localization. We generated a C-terminal fusion of superfolder GFP (sGFP) to *P. aeruginosa* DprE1, co-expressed the fusion with control protein PilE-mCherry in *E. coli* DH5α and in Pa5196 Δ*dprE1*, and examined localization of the fusions by fluorescence microscopy. PilE is a T4P minor pilin that localizes to the inner membrane with its C-terminus in the periplasm, resulting in circumferential fluorescence (23). While PilE-mCherry showed the expected pattern of fluorescence, the *P. aeruginosa* DprE1-sGFP fusion showed diffuse localization in both *E. coli* and Pa5196 strains, consistent with cytoplasmic expression (Fig. 7A **and** B). This result, coupled with the BACTH interaction data, suggests that precursor biosynthesis is likely mediated by a cytoplasmic biosynthetic complex.

**Figure 7.**
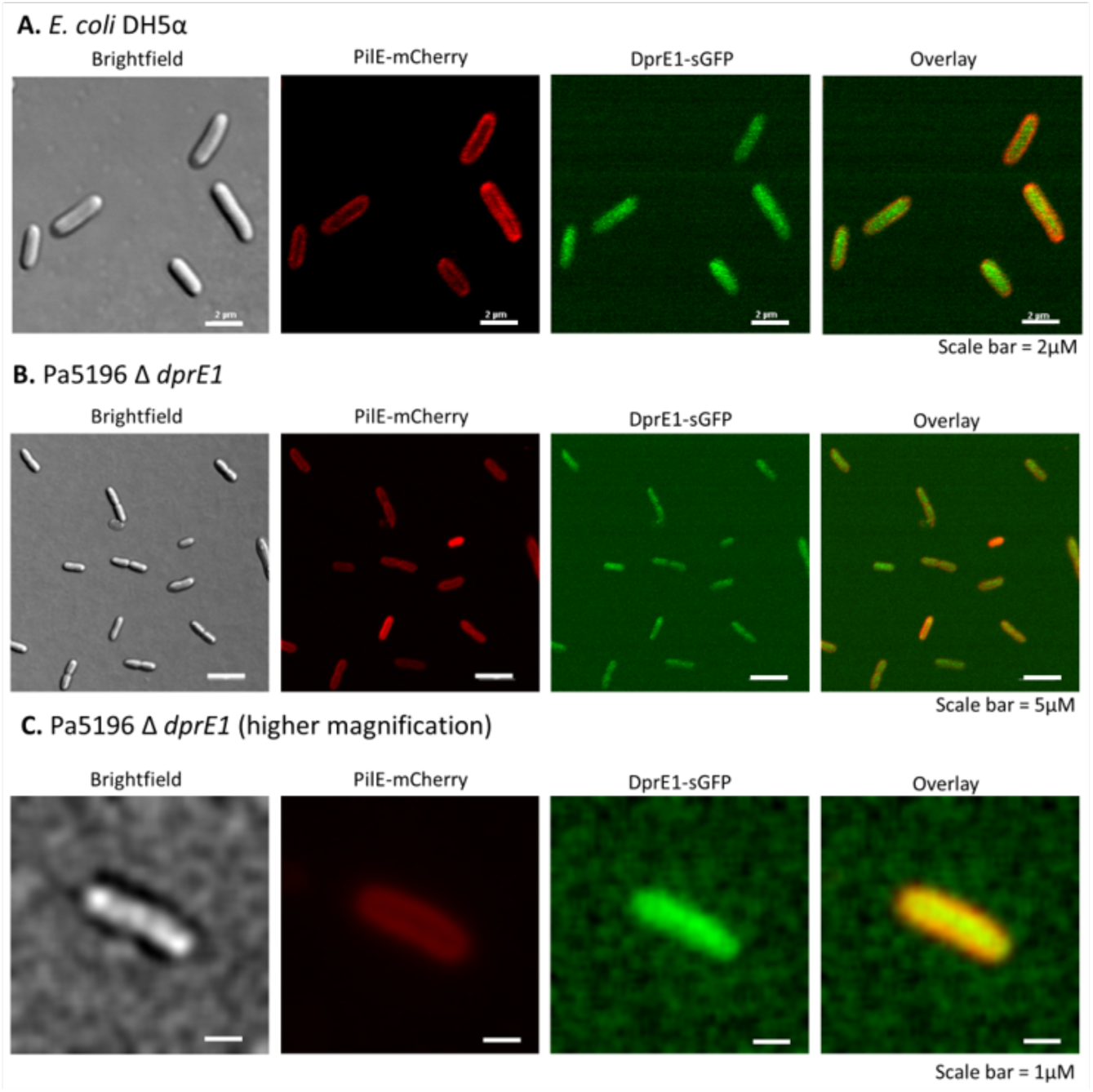
DprE1 (6248) from *P. aeruginosa* Pa5196 is expressed in the cytoplasm of *E. coli* (A) and Pa5196 Δ *dprE1* (B) strains. A higher magnification image of Pa5196 Δ *dprE1* strain is shown (C). Fluorescent microscopy analysis of DprE1-sGFP localization shows expression in the cytoplasm (green) while PilE-mCherry is expressed in the periplasm (red) of the cell. The higher magnification image of Pa5196 Δ *dprE1* strain was imaged using an EVOF FL Auto microscope through a Plan Apo 60X (NA=1.40) oil objective in the McMaster Biophotonics Facility, as described on experimental procedure session.

## DISCUSSION

TfpW is an unusual protein with features of both glycosyltransferases and oligosaccharyltransferases. TfpW and its homologues have conserved DDX (5) and proline-rich motifs (YX_6_PFX_7_P) (Fig. 1B **and** 2B). The proline-rich motif is found in membrane-bound polysaccharide polymerases including EmbA/B/C (*M. tuberculosis*), ExoP (*Sinorhizobium meliloti*), CtrB (*N. meningitidis*), WzzB (*Salmonella typhimurium*) and WzzE (*E. coli*), and regulates polysaccharide chain length (16,17). W627L and W627L/P635S/P641S mutations in the proline-rich motif of EmbC from *M. smegmatis* resulted in LAM polymers with shorter arabinan chains, confirming a role in chain-length regulation (16). The periplasmic location of TfpW’s DDX motif and TPR domains were supported by our topology studies (Fig. 3A) but fusions to other predicted periplasmic loops gave indeterminate signals. Alexeyev and Winkler reported that blue and purple colonies can result from PhoA/LacZα fusions to periplasmic domains and adjacent transmembrane regions (24). Structural studies will be necessary to validate the conformation of TfpW, particularly the central region of the protein.

The D-Ara*f* biosynthetic enzymes minimally required for modification of PilA_5196_ in the PAO1 background were 6246, DprE1 and DprE2 (9), while TfpX was dispensable (Fig. 4A). These data suggest that TfpW alone may polymerize α-1,5-D-Ara*f* glycans and transfer them to pilins. This finding is consistent with the lack of TfpX homologues in most species that encode TfpW homologues. However, TfpX was required for pilin glycosylation in *E. coli* (Fig. 4B **and** C), possibly explaining why it has been retained in some species. TfpX is a protein of unknown function with ∼50% amino acid sequence similarity to TfpY and TfpZ from group III and V strains of *P. aeruginosa*, respectively (25). Loss of those components reduces pilus assembly in an unknown manner.

GtrA-like protein 6247 shares of the properties of Rv3789 from *M. tuberculosis* despite lack of sequence similarity. GtrA family members are involved in the synthesis of cell surface polysaccharides (18). Rv3789 was initially proposed to be a flippase responsible for translocation of DPA across the cytoplasmic membrane, due to a reduction in the arabinose content of both AG and LAM and accumulation of DPA in *M. smegmatis* mutants that were restored upon complementation with *E. coli arnE/F* (19). ArnE and ArnF form a heterodimer to transport undecaprenyl phosphate-α-4-amino-4-deoxy-L-arabinose (L-Ara4N) to the outer surface of the inner membrane (26). Then, ArnT, an integral membrane glycosyltransferase, modifies the lipid A moiety of LPS with L-Ara4N, reducing its negative charge to minimize interaction with polymyxin and cationic antimicrobial peptides (27). The L-Ara4N pathway is a major contributor to antibiotic resistance in many species, including *P. aeruginosa* (7).

Based on the ability of ArnE/F to replace Rv3789 from *M. tuberculosis (19)*, they have relaxed substrate specificities regarding both oligosaccharide and lipid moieties *(26).* In *P. aeruginosa*, deletion of *arnE/F* had no effect on pilin glycosylation when AWX and 6245-6251 were expressed (Fig. 5A **and** B). However, in PAO1 Δ*arnE/F* loss of 6247 expression decreased pilin glycosylation, suggesting that these genes may have a role when 6247 is absent. Since some pilin glycosylation was still observed when both ArnE/F and 6247 were missing (Fig. 5B), it is unlikely that these proteins are responsible for D-Ara*f* translocation.

Interaction of Rv3789 with arabinosyltransferase AftA was suggested to be consistent with Rv3789 acting as a hub for other enzymes involved in DPA biosynthesis (18). Kolly et al. argued that the structural model of Rv3789 was inconsistent with a flippase activity, while its positive charge at neutral pH (pI=10.4) could facilitate the recruitment of partner proteins (18). Deletion of the Rv3789 homologue 6247 in PAO1 decreased the abundance of highly glycosylated species (9) but 6247 expression was not essential for pilin glycosylation. Our BACTH data supports a role for 6247 as a potential hub protein that interacts with TfpW, TfpX, DprE1 and DprE2 to enable efficient pilin glycosylation (Fig. 6). Our DprE1 localization data suggest that the precursor is synthesized in the cytoplasm, although no specific translocase/flippase candidate was identified. From our data, we propose a model of Pa5196 pilin glycosylation, in which a dimer of 6247 interacts with cytoplasmic DprE1 and DprE2 enzymes (Fig. 8). 6247 also recruits TfpX and TfpW to form a glycosylation complex (Fig. 8). With its limited similarity to glycan flippases, TfpW might be a trifunctional enzyme that translocates the lipid-linked precursor to the periplasmic face of the membrane, polymerizes it into α-1,5-linked-D-arabinofuranose polymers via its periplasmic DDX motif with lengths dictated by the proline-rich motif, then transfers the polymers to multiple Ser and Thr residues of PilA (Fig. 8). TfpX and the GtrA-like 6247 protein may be necessary to stabilize the pilin glycosylation complex in *E. coli*. This latter result suggests that ArnE/F may compensate for 6247 in *P. aeruginosa*, while the *E. coli* homologues might be unable to do so.

**Figure 8.**
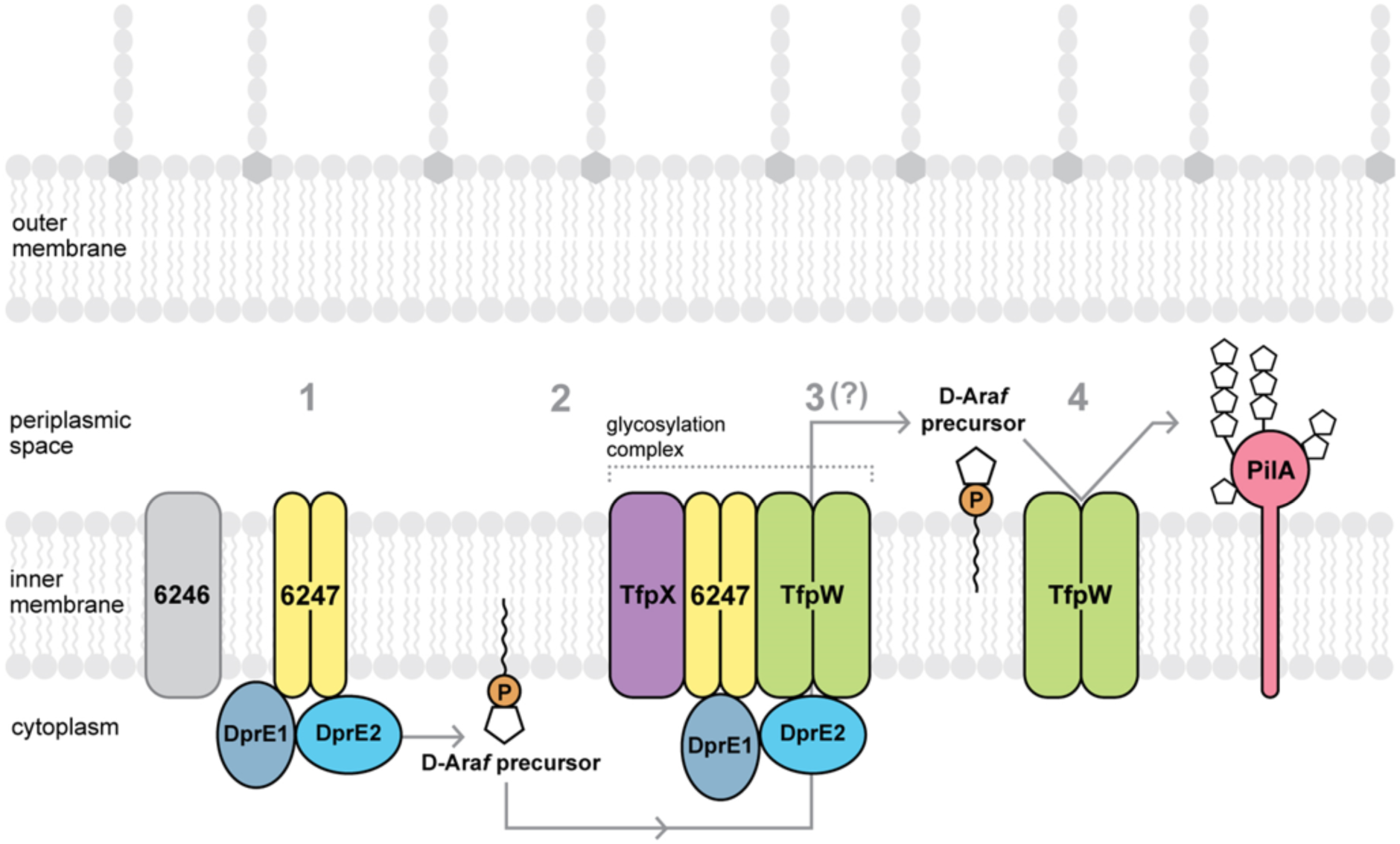
Proposed model of Pa5196 pilin glycosylation. A dimer of the GtrA-like protein 6247 interacts with DprE1 (6248) and DprE2 (6249), to form the D-Ara*f* precursor (step 1). 6247 recruits TfpX and TfpW dimers that interact to form the glycosylation complex (step 2). The D-Ara*f* precursor might be translocated from cytoplasm to periplasm by TfpW (step 3). TfpW polymerizes D-Ara*f* precursors into α-1,5-linked-D-arabinofuranose polymers, and transfers them to multiple Ser and Thr residues of PilA (step 4). Other members of the glycosylation complex, TfpX, 6247, DprE1 and DprE2 were not included on step 4 for concision. 6246, a DPPR synthetase, is essential for pilin glycosylation. Further studies are necessary to investigate whether 6246 interacts with other proteins involved in D-Ara*f* biosynthesis. ArnE/F may replace 6247 function acting as a hub protein and enabling formation of the glycosylation complex.

In summary, these data suggest that GtrA-like protein 6247 acts as a hub protein and forms a glycosylation complex with TfpW, TfpX and proteins involved in D-Ara*f* biosynthesis for efficient pilin glycosylation. TfpW may be the first example of a trifunctional flippase, polymerase and glycosyltransferase. Further studies are necessary to investigate the function of TfpW as a D-Ara*f* translocase.

## EXPERIMENTAL PROCEDURES

### Bacterial strains and growth conditions

Bacterial strains and plasmids used in this work are listed in Table 1, and oligonucleotide primer sequences are listed in Table 2. *E. coli* and *P. aeruginosa* were grown at 37 °C in Luria-Bertani (LB) broth or on LB agar plates (1.5% agar) supplemented with the following antibiotics when required: 100 µg ml^−1^ ampicillin, 50 µg ml^−1^ kanamycin, 15 µg ml^−1^ gentamicin for *E. coli* and 30 µg ml^−1^ for *P. aeruginosa*. L-arabinose was added to induce expression from pBADGr *ara* promoter at concentrations of 0.2% for complementation of *E. coli* and 0.02% or 0.05% for *P. aeruginosa*. Plasmids were transformed by heat shock into *E. coli* competent cells, and by electroporation into *P. aeruginosa* cells. Standard PCR and cloning techniques were used to generate complementation constructs as listed in Table 1. Restriction, DNA polymerase and DNA ligase enzymes were from Thermo Scientific and were used according to manufacturer’s recommendation. All constructs were verified by DNA sequencing (MOBIX McMaster University). Bacterial strains were stored at −80 °C in LB containing 15% glycerol.

**Table 1.**
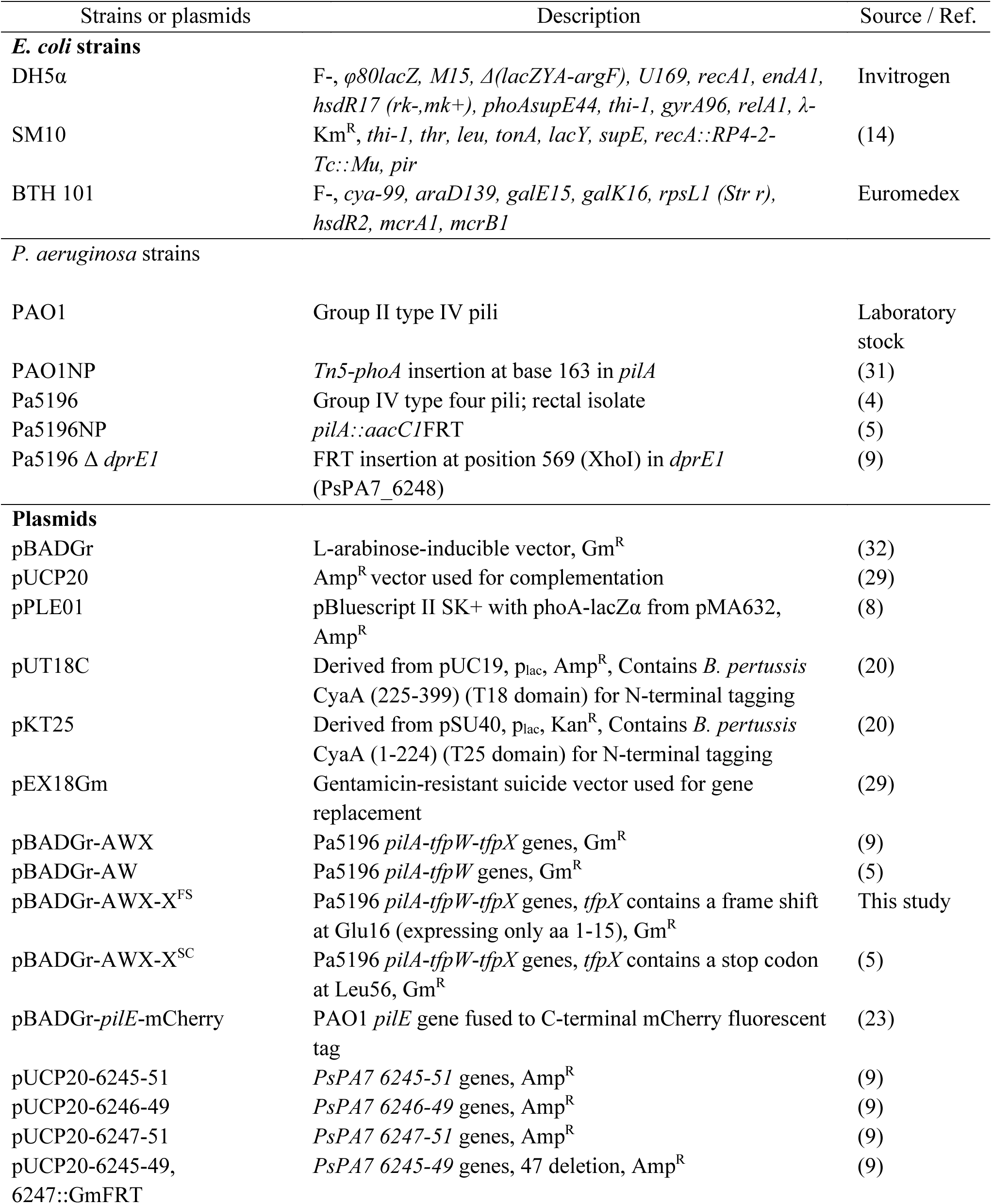
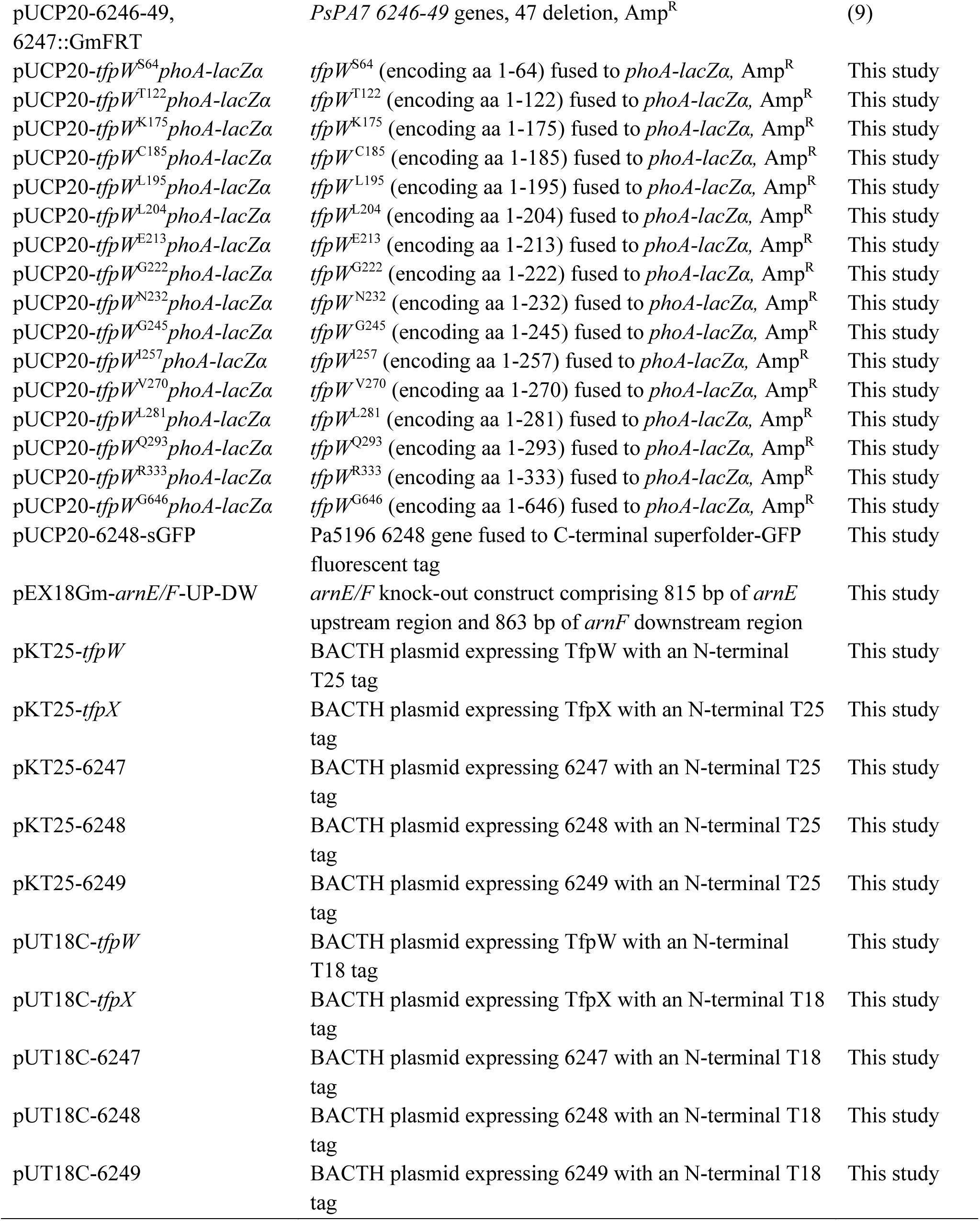
Bacterial strains and plasmids.

**Table 2.**
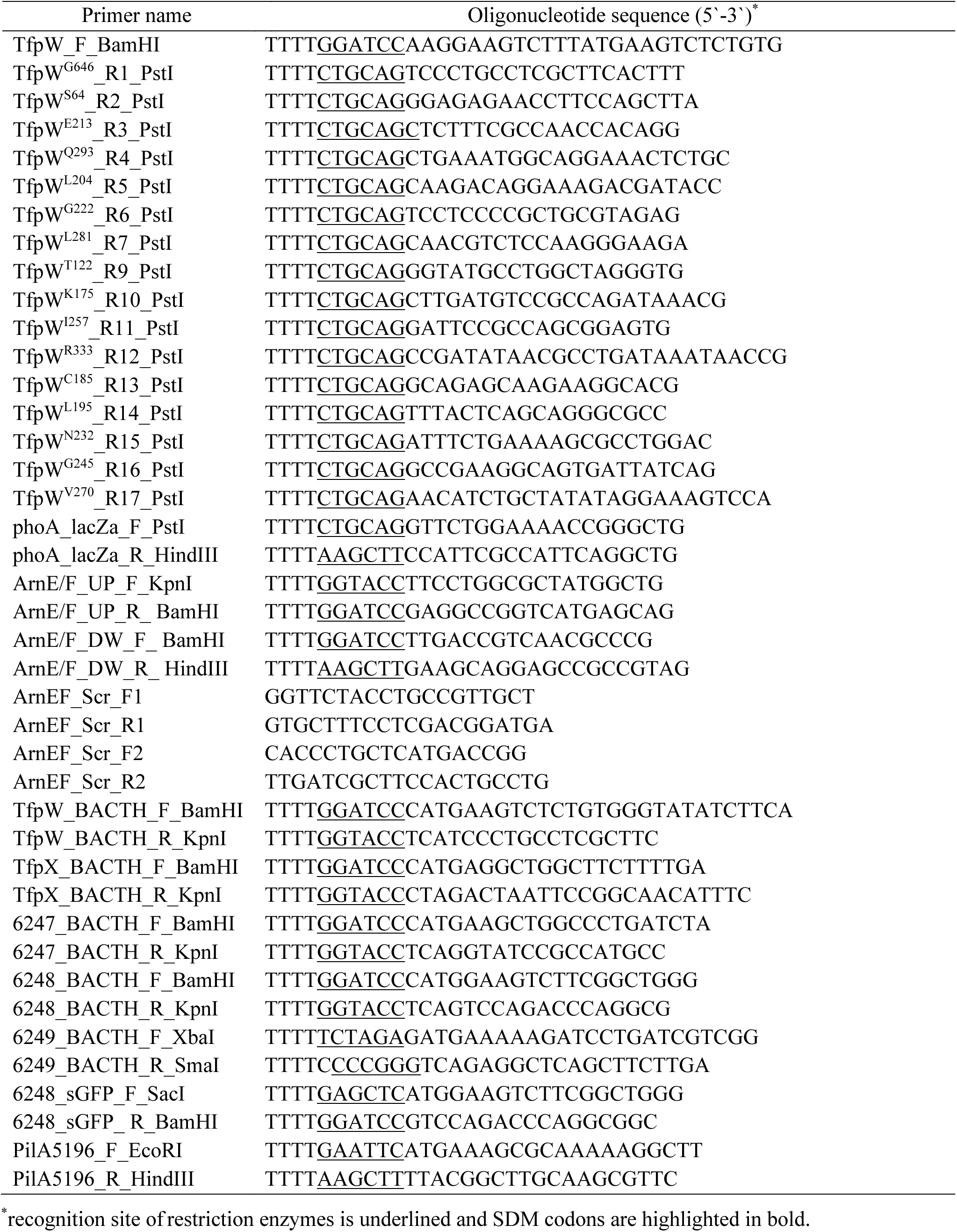
Oligonucleotide primer sequences.

### Bioinformatic analysis

TfpW putative homologs were identified through a BLASTP search (https://blast.ncbi.nlm.nih.gov/). Amino acid sequence alignments were performed using ClustalW and GONNET algorithm (https://npsa-prabi.ibcp.fr). Topology predictions were done using CCTOP (Constrained Consensus Topology prediction server) that uses 10 different topology prediction methods and incorporates information available in the PDBTM, TOPDB and TOPDOM databases using hidden Markov model (28). Structure predictions were carried out using Phyre2 web portal for protein modeling, prediction and analysis (15).

### Construction of PhoA-LacZα fusions and topology studies

The *phoA-lacZα* reporter sequence was amplified from pPLE01 (8) and cloned into pUCP20 using PstI and HindIII enzyme restriction sites. Targeted fusions for *tfpW* gene were created by amplifying various 3’ truncation positions from *P. aeruginosa* Pa5196 genomic DNA and cloning upstream of *phoA-lacZα* in pUCP20 using BamHI and PstI sites. Constructions were transformed into *E. coli* DH5α competent cells (α-complementing host strain) and plated on LB agar containing ampicillin (100 µg ml^−1^), isopropyl-β-D-thiogalactopyranoside (IPTG, 1 mM), 5-bromo-4-chloro-3-indolyl phosphate (BCIP, 80 µg ml^−1^), and 6-chloro-3-indolyl-β-D-galactoside (Red-Gal, 100 µg ml^−1^). Plates were incubated at 37 °C overnight. Colony colors represent the cellular localization of a given truncation: blue, periplasm (AP activity); red, cytoplasm (BG activity); and purple, transmembrane (combination of AP and BG activities). Three independent experiments were performed.

### Preparation of whole cell lysates

*E. coli* and *P. aeruginosa* strains were grown on LB agar plates overnight at 37 °C. Cells were scraped and resuspended in 1X PBS to an OD_600_ of 0.5 for *E. coli* or 0.1 for *P. aeruginosa* strains, and 1 ml was centrifuged at 13400 x *g* for 3 min. The cell pellet was resuspended in 100 µl of 1X SDS-PAGE loading buffer (50 mM Tris HCl pH 6.8, 100 mM DTT, 2% (w/v) SDS, 0.1% (w/v) bromophenol blue and 10% glycerol).

### SDS-PAGE and Western blot analyses

Samples were boiled at 100 °C for 10 min and separated on 15% SDS-PAGE at 150 V for 1 h. Proteins were transferred to nitrocellulose membrane for 1 h at 225 mA. Membranes were blocked with 5% skim milk powder in 1X PBS shaking for 1 h at room temperature and were incubated with primary antibodies for 16 h at room temperature. Anti-PilA_IV_ (rabbit number 286, 1:5000 dilution in 1X PBS) was used for detection of pilins. Membranes were washed three times with 1X PBS for 5 min and incubated with goat anti-rabbit IgG-alkaline phosphatase-conjugate secondary antibody (Bio-Rad, 1:3000 dilution) for 1 h at room temperature. Membranes were washed three times with 1X PBS for 5 min and developed with nitro blue tetrazolium/5-bromo-4-chloro-3-indolyl phosphate in alkaline phosphatase buffer (Bio-Rad).

### Generation of arnE/F mutant of PAO1

Primers were designed to amplify approximately 800 bp fragments corresponding to regions upstream and downstream of the *arnE/F* genes (Table 2). After digestion with restriction enzymes, upstream (KpnI and BamHI) and downstream (BamHI and HindIII) fragments were sequentially cloned into the suicide vector pEX18Gm (29). *E. coli* SM10 was transformed with the constructs and they were transferred to PAO1 by biparental mating as described previously (29). PAO1 recombinant strains were selected on *Pseudomonas* Isolation Agar (PIA, BD) containing 100 µg ml^−1^ gentamicin. PAO1 mutants were selected on LB agar containing 5% sucrose and were gentamicin-sensitive. Deletion of *arnE/F* genes was verified by PCR using screening primers (Table 2).

### Bacterial adenylate cyclase two-hybrid assay

The *tfpW*, *tfpX*, 6247, 6248 and 6249 genes were amplified by PCR from Pa5196 genomic DNA and cloned into both pUT18C and pKT25 plasmids using restriction enzymes. The pUT18C and pKT25 plasmids express T18 and T25 domains of adenylate cyclase, respectively. Competent *E. coli* BTH 101 cells were co-transformed with derivatives of both plasmids and plated on MacConkey agar supplemented with 1% maltose, 0.5 mM IPTG, 100 µg ml^−1^ ampicillin, 50 µg ml^−1^ kanamycin, and on LB agar supplemented with 40 µg ml^−1^ 5-bromo-4-chloro-3-indolyl-β-D-galactopyranoside (X-Gal), 0.5 mM IPTG, 100 µg ml^−1^ ampicillin, 50 µg ml^−1^ kanamycin, and incubated at 25 °C for 24-48 h. Three independent experiments were performed.

### Fluorescence microscopy

The *dprE1* (6248) gene fused to sGFP C-terminal tag was cloned into pUCP20 and transformed in DH5α *E. coli* and in Pa5196 Δ *dprE1* strains. Strains were also transformed with pBADGr-*pilE*-mCherry construct (23) and plate into LB agar containing 15 µg ml^−1^ gentamicin and 100 µg ml^−1^ ampicillin (*E. coli*) or 30 µg ml^−1^ gentamicin and 200 µg ml^−1^ carbenicillin (*P. aeruginosa*) containing 0.02% L-arabinose. Bacteria were spotted on a 1% agarose pad supplemented with 0.02% L-arabinose on a microscope slide and incubated at 37°C for 2-4 hours. The agarose pad was mounted with a glass coverslip directly prior to imaging. Cells were imaged using brightfield and fluorescence microscopy on a Nikon A1 confocal microscope through a Plan Apo 60X (NA=1.40) oil objective. Image acquisition was done using Nikon NIS-Elements Advanced Research (Version 5.11.01 64-bit) software. Where indicated, cells were imaged using an EVOF FL Auto microscope through a Plan Apo 60X (NA=1.40) oil objective in the McMaster Biophotonics Facility. TexasRed and YFP LED filters were used for fluorescence imaging, and transmitted white light was captured with a monochrome camera. Image analysis was completed using ImageJ 1.52K containing the MicrobeJ plugin for fluorescence channel alignment (30).

## Data availability

All data are contained within the manuscript.

## Acknowledgements

This work was funded by grants from Glyconet and the Boris Family Foundation.

